# Structural conservation of the gabapentinoid binding site in human and *Caenorhabditis elegans* α2δ subunits: a docking and molecular dynamics perspective

**DOI:** 10.64898/2026.09.15.751774

**Authors:** Jabin Sultana, Lusine Mkrtchyan, Jesus D. Castaño, Jérôme R. E. del Castillo, Francis Beaudry

**Affiliations:** Département de Biomédecine Vétérinaire, Faculté de Médecine Vétérinaire, Université de Montréal, Saint-Hyacinthe, Québec, Canada; Centre Interdisciplinaire de Recherche sur le Cerveau et l’Apprentissage (CIRCA), Université de Montréal, Montréal, Québec, Canada; Department of Physiology, Biochemistry and Pharmacology, Faculty of Veterinary Medicine, Chattogram Veterinary and Animal Sciences University, Chattogram, Bangladesh; Proteomics Platform, CHU de Québec-Université de Laval Research Center, Québec, Canada

**Keywords:** Gabapentin, Pregabalin, voltage-gated calcium channels, *in silico* analysis, molecular dynamics

## Abstract

Pain is a global health burden requiring effective therapeutic strategies. Gabapentin (GBP) and pregabalin (PGB), used for neuropathic pain, act primarily through α2δ subunits of voltage-gated calcium channels. *Caenorhabditis elegans* expresses UNC-36, an ortholog of mammalian α2δ proteins involved in calcium-channel function and nocifensive behavior. However, whether UNC-36 retains the features essential for gabapentinoid recognition remains unclear. We compared human α2δ-1 and UNC-36 using sequence and structural analyses, docking, one 500-ns molecular dynamics trajectory per complex, distance-based contact profiling, principal component and free-energy landscape analyses, and MM/GBSA calculations. The modeled proteins showed Cα correspondence within a selected 13-residue pocket core (RMSD 0.53 Å), whereas the complete 14-residue mapping yielded 9.07 Å. All four trajectories showed recurrent ligand–protein proximity contacts with residue-level patterns. Aromatic-residue contacts accounted for a larger fraction of contact records in human trajectories, whereas basic-residue proximity was more frequent in UNC-36 trajectories, including recurrent Arg501 and Arg503 contacts. Human α2δ-1– PGB showed the lowest ligand RMSD, while α2δ-1–GBP showed greater positional variation. In UNC-36, PGB deviated more from its initial pose than GBP. The first two principal components accounted for more conformational variance in UNC-36 complexes than in human complexes. MM/GBSA estimates, without an explicit conformational-entropy contribution, were more negative for PGB than GBP in human trajectories and for GBP than PGB in UNC-36 trajectories. These simulations suggest that selected pocket-core correspondence coexists with differences in modeled contacts, sampled motions, and estimates. They provide hypotheses for investigating gabapentinoid recognition by UNC-36 and its pharmacological relevance.

## 1 Introduction

Chronic pain is one of the major causes of disability and it remains difficult to treat because the mechanisms that initiate and sustain pain vary among patients and disease states (1–4). This heterogeneity is particularly evident in neuropathic pain, for which currently available therapeutics often provide incomplete or partial relief (5). Among commonly prescribed non-opioid analgesics, gabapentin (GBP) and pregabalin (PGB), collectively termed gabapentinoids, are widely used for the management of neuropathic pain and partial-onset epilepsy (6). Despite their structural similarity to γ-aminobutyric acid (GABA), neither compound acts primarily through classical GABA receptors. Instead, they bind with high affinity to the α2δ auxiliary subunits (α2δ-1 and α2δ-2) of voltage-gated calcium channels (VGCCs) (7,8).

The α2δ subunits regulate several aspects of voltage-gated calcium channel biology and function, including channel trafficking, membrane expression, and calcium-dependent neurotransmitter release (9,10). Structural studies have localized the gabapentinoid-binding site of mammalian CaVα2δ-1 to the first dCache1 domain, an extracellular ligand-binding module that contains an evolutionarily conserved amino acid–binding pocket. Gabapentinoids bind within this pocket, where they are enclosed by a network of charged, polar, and aromatic residues (8,11). Earlier mutagenesis studies identified Arg217 as a conserved residue within this region that is critical for gabapentin binding (12). Following binding to α2δ, gabapentinoids can interfere with α2δ-dependent trafficking and surface delivery of voltage-gated calcium-channel complexes, thereby reducing presynaptic VGCC channel availability and activity-dependent neurotransmitter release (8–10). These effects can dampen enhanced excitatory transmission within sensitized nociceptive pathways and contribute to the analgesic actions of GBP and PGB.

*Caenorhabditis elegans* (*C. elegans*) is a useful whole-animal system for studying nociceptive signaling because of its well-characterized nervous system, genetic accessibility, short life cycle, and suitability for relatively high-throughput experimental approaches (13,14). Moreover, *C. elegans* exhibits reproducible thermal avoidance responses to noxious heat that make it a useful behavioral model for studying nociception and antinociceptive pharmacology (15). In *C. elegans*, voltage-gated calcium channels (VGCCs) play a central role in sensory signaling by allowing Ca²⁺ entry following membrane depolarization (16–18). This calcium influx helps the neuronal activity involved in detecting and responding to noxious stimuli. The core VGCC structure is evolutionarily conserved in *C. elegans*, although the pore-forming α1 subunits correspond to specific channel classes. EGL-19 is the CaV1/L-type α1 subunit, whereas UNC-2 is the CaV2-type presynaptic α1 subunit; UNC-36 functions as the principal neuronal α2δ auxiliary subunit, and CCB-1 represents a conserved β auxiliary subunit (19,20).

Our previous work has shown that disruption of *unc-2* or *unc-36* impairs avoidance of noxious heat, supporting the role of VGCC signaling in thermonociceptive behavior (18). In the same study, we showed that GBP and PGB significantly reduced thermal avoidance in a concentration-dependent manner, while proteomic analysis identified treatment-associated changes in proteins and signaling pathways related to VGCC function and nociceptive processing (18). Although both UNC-2 and UNC-36 contribute to VGCC-mediated nocifensive behavior, UNC-36 is especially relevant to gabapentinoid pharmacology because it is the *C. elegans* homolog of the mammalian α2δ auxiliary subunits targeted by these drugs. Therefore, the present study focuses on UNC-36 to determine whether it preserves the structural and molecular features required for gabapentinoid recognition. This work builds on a broader series of studies from our laboratory using *C. elegans* thermal avoidance to investigate analgesic pharmacology. Our team showed that capsaicin and related vanilloids modify nocifensive responses through mechanisms involving the TRPV-like channels in *C. elegans* OSM-9 and OCR-2 (21–24). Recently, molecular dynamics simulations comparing capsaicin recognition by mammalian TRPV1 and the *C. elegans* TRPV-related channels OSM-9 and OCR-2 indicated that the overall ligand-recognition architecture can be preserved despite substantial divergence in the residue-level interactions that stabilize the ligand (25). This distinction between conservation of binding-site architecture and conservation of the specific molecular contacts within it raises the question of how closely UNC-36 recapitulates the gabapentinoid-binding determinants of mammalian α2δ proteins.

Earlier electrophysiological work indicates that functional conservation does not necessarily imply that UNC-36 engages gabapentinoids in the same manner as mammalian α2δ proteins (19). That study also showed that UNC-36 contributes to calcium influx in cultured *C. elegans* mechanosensory neurons and identified differences in several residues previously reported for gabapentin recognition in mammalian α2δ proteins. This is particularly relevant because the gabapentinoid-binding region of UNC-36 has not yet been structurally characterized. In mammalian α2δ-1, GBP binds within the amino acid–binding pocket of the first dCache1 domain described above, and this pocket is formed by a spatially clustered set of charged, polar, and aromatic residues that surround the ligand and contribute to its recognition (8,11). These structurally characterized residues therefore provided the reference for identifying and aligning the corresponding region of UNC-36 and for defining the candidate docking site. However, it remains unclear whether the corresponding region of UNC-36 retains a similar physicochemical environment and steric complementarity required for GBP and PGB recognition. Therefore, a direct structural comparison is necessary to determine whether conservation of α2δ-associated function is accompanied by conservation of the molecular environment responsible for gabapentinoid recognition. Molecular docking and molecular dynamics (MD) simulations offer complementary means of addressing this question: docking identifies plausible binding orientations, while MD samples positional changes and ligand-protein contact patterns over time (26,27). Combined with residue-level contact profiling, conformational analysis, and MM/GBSA binding-energy estimates calculations, these approaches allow the comparison of conserved and divergent features of ligand recognition across homologous proteins (28). Establishing the extent to which UNC-36 reproduces the gabapentinoid-binding determinants of human α2δ proteins is therefore important for interpreting *C. elegans* pharmacological findings in a translational context, because target correspondence is one factor determining how observations in the nematode can inform mammalian pharmacology.

In the present study, we investigated the molecular recognition of GBP and PGB by human CaVα2δ-1 and *C. elegans* UNC-36. Sequence and structural comparisons of the gabapentinoid-binding region were combined with molecular docking and one 500-ns MD trajectory for each of the human α2δ-1–GBP, human α2δ-1–PGB, UNC-36–GBP, and UNC-36–PGB complexes. The resulting trajectories were examined using structural stability metrics, time-resolved contact profiling, cumulative residue contact-class counts, principal component analysis, free-energy landscape analysis, and MM/GBSA binding-energy calculations. We hypothesized that UNC-36 retains a structurally corresponding core within the candidate gabapentinoid-binding region related to the mammalian α2δ-1 dCache1 site, but that species-specific sequence differences modify the residue-level interactions through which GBP and PGB are accommodated.

The four final production systems are referred to as HS+GBP (human α2δ-1 with gabapentin), HS+PGB (human α2δ-1 with pregabalin), CE+GBP (UNC-36 with gabapentin), and CE+PGB (UNC-36 with pregabalin) throughout the manuscript.

## 2 Results

### 2.1. Sequence and structural comparisons identify a selected conserved pocket core

The full-length human α2δ-1 and *C. elegans* UNC-36 sequences were substantially divergent, but conservation was concentrated at several positions associated with the modeled ligand-binding environment. Human CACNA2D1 and UNC-36 were aligned before docking to identify the corresponding gabapentinoid-binding region. Following the MD simulations, residues with recurrent proximity contacts to GBP or PGB were mapped onto the final MAFFT alignment to assess their conservation between the two proteins. Figure 1A shows identical residues at Tyr217/Tyr215, Trp223/Trp221, Trp243/Trp240, Tyr450/Tyr500, and Leu456/Leu506, together with the conservative aromatic substitutions Trp205/Phe203 and Tyr236/Phe233. In each pair, the first residue refers to human α2δ-1 and the second to UNC-36. This arrangement is consistent with the modular architecture of α2δ proteins, in which residues contributing to the three-dimensional gabapentinoid-recognition site can be brought together from separated sequence regions within the folded extracellular domain (11).

**Figure 1.**
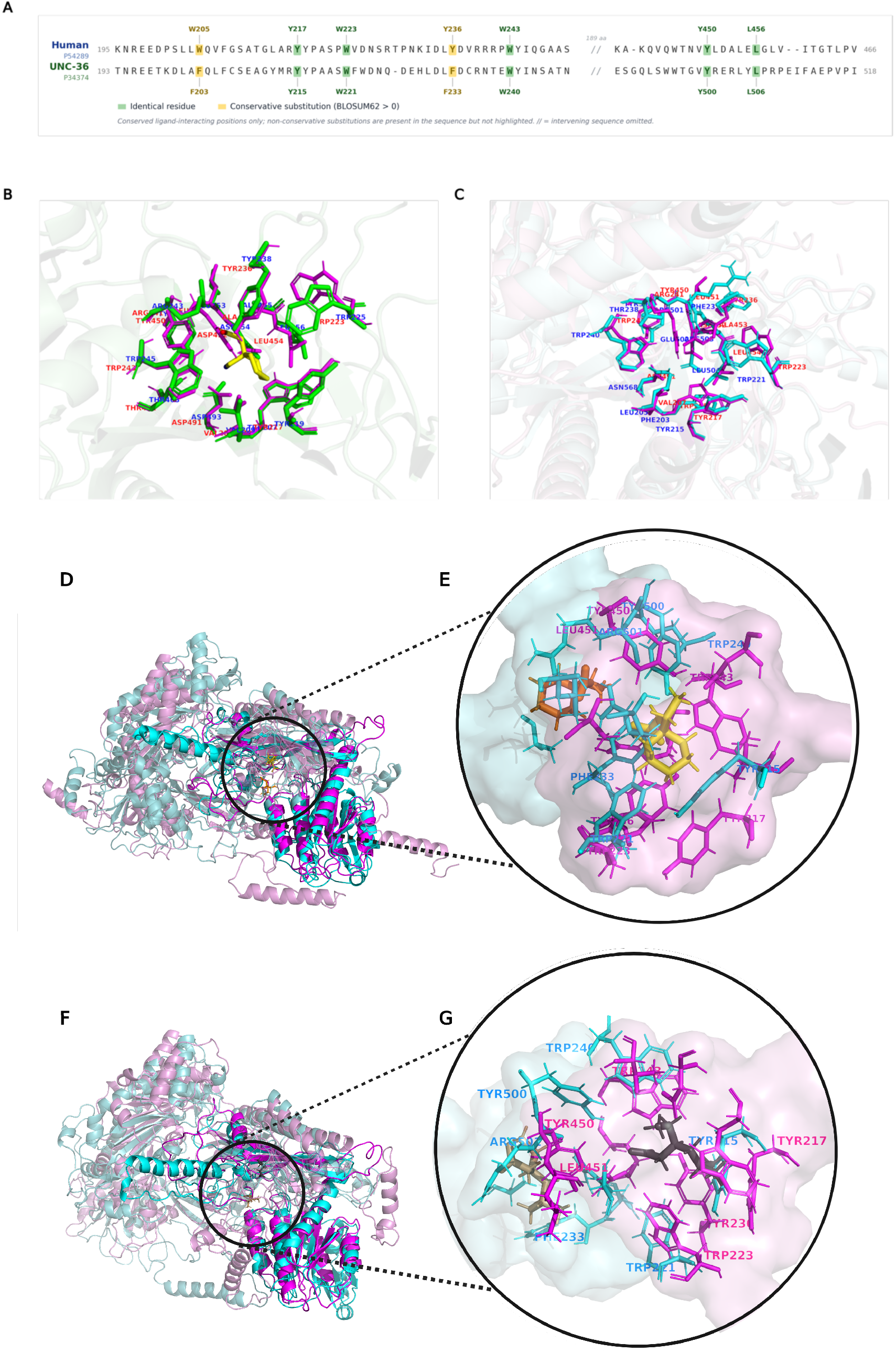
Sequence and structural comparison of the gabapentinoid-binding region in mammalian α2δ-1 and *C. elegans* UNC-36. **(A)** MAFFT alignment of human CACNA2D1 and UNC-36; identical residues are shown in green and conservative substitutions in yellow. **(B)** Superposition of the experimental rabbit CaVα2δ-1 gabapentin-binding pocket from PDB 8FD7 (green) and the corresponding predicted human pocket (magenta); GBP is shown in yellow. **(C)** Superposition of the selected 13-residue Cα pocket cores of human α2δ-1 (magenta) and UNC-36 (cyan), excluding the Thr461/Ala513 pair. **(D, F)** Full-length GBP and PGB systems, respectively, after domain-restricted superposition, with human α2δ-1 shown in magenta and UNC-36 in pale cyan. **(E, G)** Enlarged GBP- and PGB-centered pocket views from the final MD snapshots, with pocket surfaces defined by residues within 5 Å of the ligand and selected residues shown as sticks; human and UNC-36 ligands are shown in yellow/orange for GBP and dark gray/light brown for PGB, respectively.

To determine whether this sequence-level conservation corresponded to the experimentally established mammalian gabapentinoid-binding site, we compared the models with the cryo-EM structure of gabapentin-bound rabbit CaVα2δ-1 (UniProt P13806) within the CaV1.2 channel complex (PDB 8FD7). Rabbit and human α2δ-1 are highly conserved across the dCache1 gabapentin-binding region, including the identified ligand-contacting positions (11) (Figure 1B). To our knowledge, no cryo-EM structure of human α2δ-1 directly bound to gabapentin or pregabalin is currently available. Although the α2δ-1 subunit in 8FD7 is from rabbit, the original structural study showed strong conservation of the gabapentin-binding region between rabbit and human α2δ proteins, including the residues involved in ligand recognition. Therefore, 8FD7 was used as the experimental reference for defining the α2δ-1 dCache1 gabapentin-binding pocket (11). Fourteen α2δ-1 residues located within 5 Å of GBP in the experimental 8FD7 structure were identified, and the corresponding positions were determined in the human α2δ-1 (CACNA2D1) model and subsequently in UNC-36. To assess the local accuracy of the human α2δ-1 model, the Cα atoms of the 14 equivalent binding-pocket residues were superposed onto those of the experimental structure (PDB 8FD7), giving an RMSD of 0.63 Å. Because this comparison is restricted to backbone Cα positions, it supports the modeled pocket backbone geometry only; it does not validate side-chain placement or provide information on ligand affinity. Sequence alignment was used to identify the corresponding positions in UNC-36. The 14 sequence-mapped residues were then structurally superimposed with the corresponding residues in the human α2δ-1 model, yielding a Cα RMSD of 9.07 Å. However, structural inspection showed that this value was strongly influenced by the human Thr461/UNC-36 Ala513 pair, whose Cα atoms were separated by approximately 36.3 Å after alignment of the surrounding pocket core. Thus, although these residues were aligned at the sequence level, they did not occupy spatially equivalent positions in the compared models. The original UNC-36 prediction assigned Ala513 a predicted local distance difference test (pLDDT) score of 44.25, indicating low local confidence. Its apparent displacement therefore cannot be reliably interpreted as a genuine structural difference between UNC-36 and human α2δ-1. Excluding this pair from the structural comparison yielded a Cα RMSD of 0.53 Å for the selected 13-residue core. This lower value describes the selected core and does not establish equivalence of the complete 14-residue region or its side-chain geometries. The displaced pair remains relevant because human Thr461 was a recurrent contact in the HS+PGB trajectory. Moreover, local comparisons further supported this structural correspondence. The first pocket-forming region, comprising human residues Trp205–Trp243 and their mapped UNC-36 counterparts, superimposed with a Cα RMSD of 0.40 Å. A second core segment, human Tyr450– Leu454 and UNC-36 Tyr500–Leu504, showed an even lower Cα RMSD of 0.30 Å. Despite the closely conserved backbone geometry, the aligned residues differed in amino-acid identity and side-chain physicochemical properties. Notably, human Leu451 and Ala453 corresponded to Arg501 and Arg503 in UNC-36, respectively, introducing two positively charged side chains into a region with closely conserved backbone geometry. Human Arg241, in contrast, corresponded to Thr238 in UNC-36, indicating that the distribution of positive charge within the pocket is reorganized rather than simply increased. Consistent with this difference, Arg501 and Arg503 were subsequently identified as recurrently contacted residues for GBP and PGB during the UNC-36 MD trajectories, as discussed later. Together, these comparisons support local backbone correspondence within the selected core while identifying a substantial displacement at one mapped position and marked differences in side-chain chemistry.

Additional analysis of pLDDT and predicted aligned error (PAE) provided context for confidence in the mapped pocket region (Table S1). Across the complete 14-residue mapping, mean pLDDT was 88.02 for human α2δ-1 and 88.03 for UNC-36, but the UNC-36 average concealed the low-confidence Ala513 position. In the selected 13-residue core, mean pLDDT was 87.56 (range 79.25–95.88) for human α2δ-1 and 91.39 (86.06–97.75) for UNC-36; mean directed off-diagonal PAE was 2.37 and 1.99 Å, respectively. Including the displaced pair gave mean PAE values of 2.34 and 3.75 Å. These results support greater confidence in the selected UNC-36 core than in Ala513 and its placement relative to that core. They do not validate the modeled binding poses or establish equivalent side-chain arrangements.

### 2.2. Backbone and ligand RMSD profiles differ among the four analyzed trajectories

Within the analyzed trajectories, protein backbone RMSD reached apparent plateaus rather than showing continuous drift over 500 ns (Figure 2A). HS+PGB showed the lowest backbone deviation, generally around 0.3-0.45 nm during much of the trajectory, whereas HS+GBP, CE+GBP, and CE+PGB exhibited higher RMSD ranges that stabilized broadly around 0.9-1.2 nm. The absence of sustained upward drift provided descriptive evidence of limited backbone drift over the observed time window. Radius-of-gyration profiles likewise remained within relatively narrow ranges for all four systems (Figure 2C), providing no evidence of progressive unfolding or loss of compactness.

**Figure 2.**
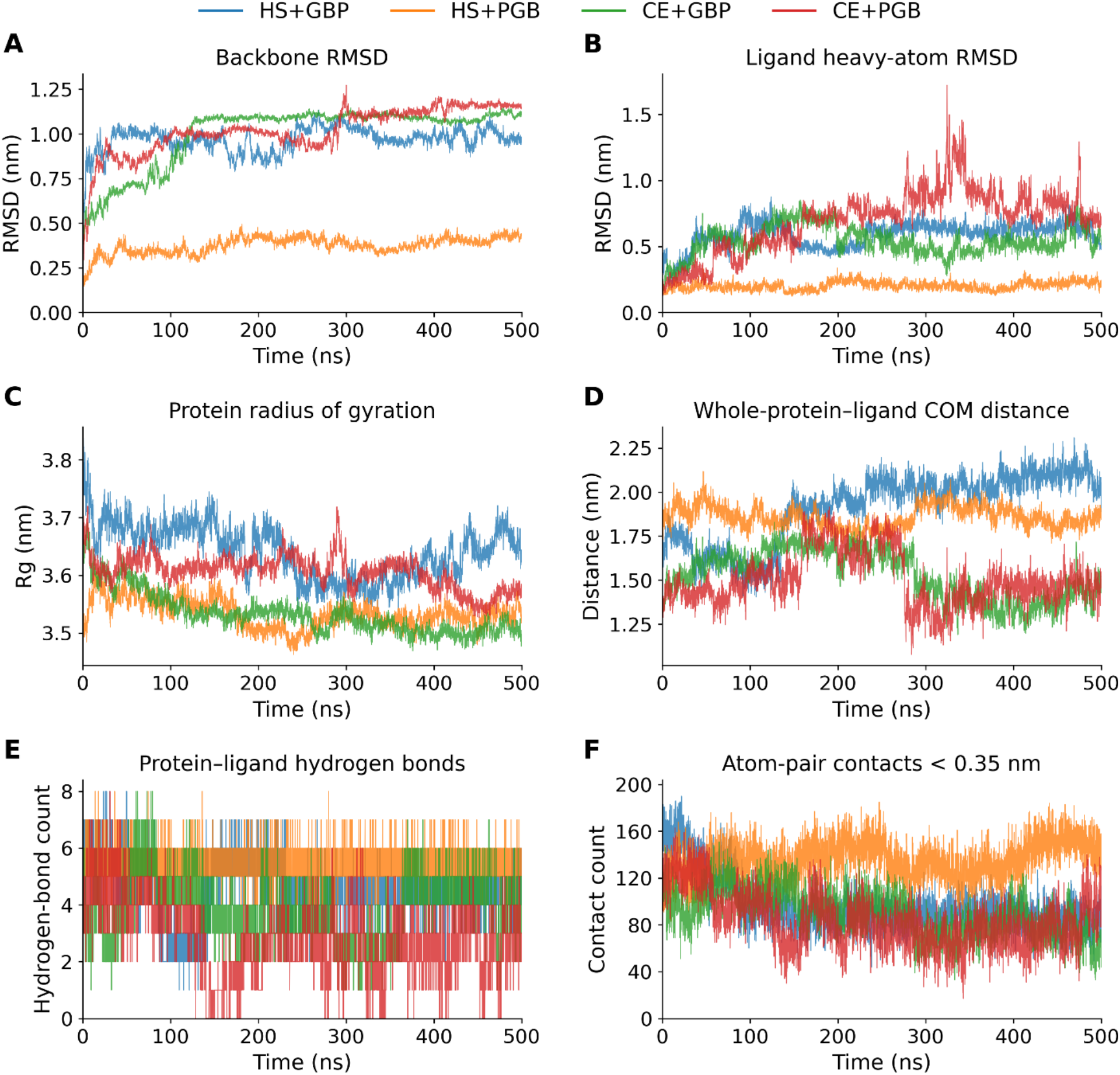
Molecular dynamics profiles of human α2δ-1 and C. elegans UNC-36 complexes with gabapentin (GBP) and pregabalin (PGB). One final 500-ns trajectory was analyzed per complex. (A) Protein backbone RMSD after backbone fitting to the production TPR reference. (B) Ligand heavy-atom RMSD from the saved corrected outputs, relative to the first frame of the protein-fitted trajectory without further fitting. (C) Protein radius of gyration. (D) Distance between the centers of mass of the whole protein and ligand; this is a global geometric measure rather than a direct measure of displacement from the binding pocket. (E) Protein–ligand hydrogen-bond counts from gmx hbond-legacy. (F) Number of protein–ligand atom pairs within 0.35 nm, including hydrogens. All 5,001 saved frames are plotted without smoothing. HS+GBP, HS+PGB, CE+GBP, and CE+PGB are shown in blue, orange, green, and red, respectively.

Ligand RMSD profiles also differed across the four trajectories (Figure 2B). In the analyzed HS+PGB trajectory, PGB showed lower deviation from its protein-aligned reference than the ligands in the other three trajectories, while GBP in both proteins showed broader positional fluctuations.

CE+PGB showed the largest transient ligand displacement, including excursions above 1.0 nm and peaks approaching 1.7 nm during the middle-to-late portion of the trajectory, followed by reorganization into a lower-RMSD region. These profiles describe ligand displacement within the sampled trajectories; they do not establish reproducible differences in ligand stability between the proteins. Hydrogen-bond and contact profiles recorded recurrent protein-ligand contacts in all four analyzed trajectories (Figure 2E-F). Within this comparison, HS+PGB showed the highest and most persistent contact counts and generally more recurrent hydrogen bonds. CE+PGB exhibited fewer and more variable hydrogen-bond interactions, while HS+GBP and CE+GBP showed intermediate hydrogen-bonding patterns. These patterns led us to examine which residues contributed to the recorded contacts in each complex.

### 2.3. Time-resolved contact profiles describe recurrent residue proximity

Residue-by-time heatmaps summarize the number of positive distance-based contact classes for each residue and frame (Figure 3; Table S4). The generic 4-Å proximity class was omitted from this display to show the remaining atom- and residue-selection classes. For readability, Figure 3 shows up to 20 highest-ranked residues with at least one nongeneric record, ranked by cumulative counts across all classes; full residue profiles were retained with the analysis outputs. These classes describe proximity and do not assign specific hydrogen bonds, aromatic interactions, or salt bridges.

**Figure 3.**
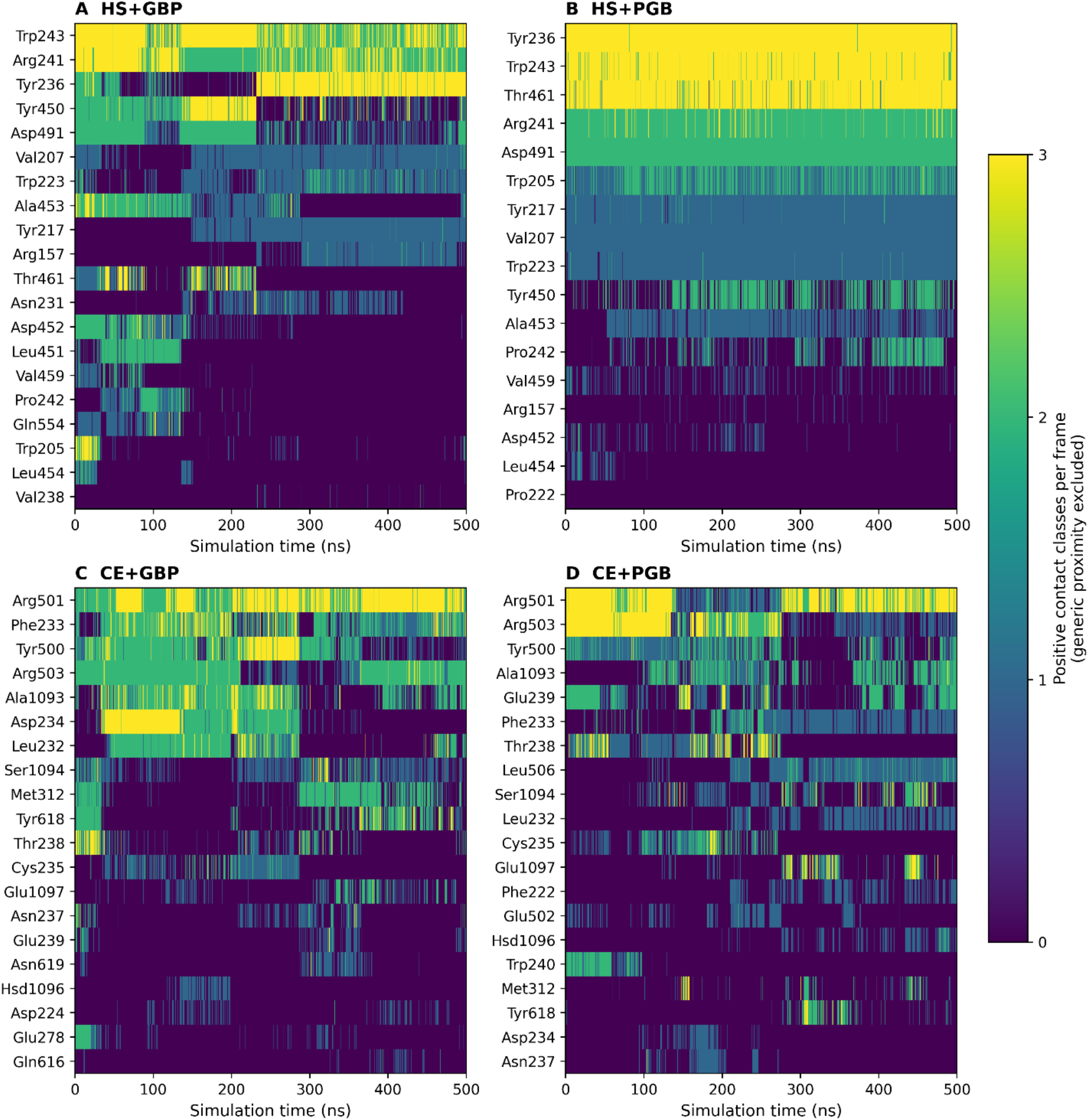
Time-resolved distance-based contact-class counts, excluding generic proximity. Panels: (A) HS+GBP, (B) HS+PGB, (C) CE+GBP, and (D) CE+PGB. Each panel shows up to 20 highest-ranked residues with at least one nongeneric record, ordered by cumulative counts across all classes. Columns correspond to 100-ps trajectory frames from 0–500 ns; color indicates the number of positive nongeneric classes for each residue and frame, with a common scale across panels. Categories use the atom selections and cutoffs in Table S4 and do not assign chemically typed interactions. Full residue profiles were retained with the analysis outputs.

In the analyzed trajectories, the ligands showed recurrent contacts involving aromatic, arginine and polar residues, with differences in contact persistence among the four complexes. In the human complexes, a common residue set was engaged repeatedly. In HS+GBP, Trp243, Arg241, Tyr236, Tyr450, Asp491, and adjacent residues contributed recurrent proximity records over the 500-ns trajectory, and HS+PGB engaged a closely related network in which Tyr236 and Trp243 remained prominent while Thr461 became particularly persistent. The two human trajectories therefore sampled overlapping sets of contacting residues, with differences in the recorded contact persistence.

The two UNC-36 trajectories showed proximity contacts with different residue distributions. CE+GBP engaged Arg501 and Arg503 together with Tyr500, Phe233, Ala1093, and residues within the 232–239 segment, and CE+PGB retained the same arginine pair as recurrently contacted residues alongside Tyr500, Phe233, Ala1093, Glu239, Thr238, and neighbouring polar residues. The same contact regions recurred in the UNC-36 trajectories, with individual proximity contacts appearing and disappearing as the ligands reoriented. These frame-wise profiles describe contacts at the 100-ps sampling interval and do not measure the lifetime of individual interactions.

### 2.4. Cumulative residue counts and contact classes differ among the sampled complexes

Ranking residues by cumulative positive contact-class records highlighted differences among the analyzed trajectories (Figure 4). In HS+GBP, Trp243, Arg241, and Tyr236 had the largest counts, followed by Tyr450 and Asp491. Each count sums binary class-presence records across frames and classes, so overlapping classes can contribute more than once per residue and frame.

**Figure 4.**
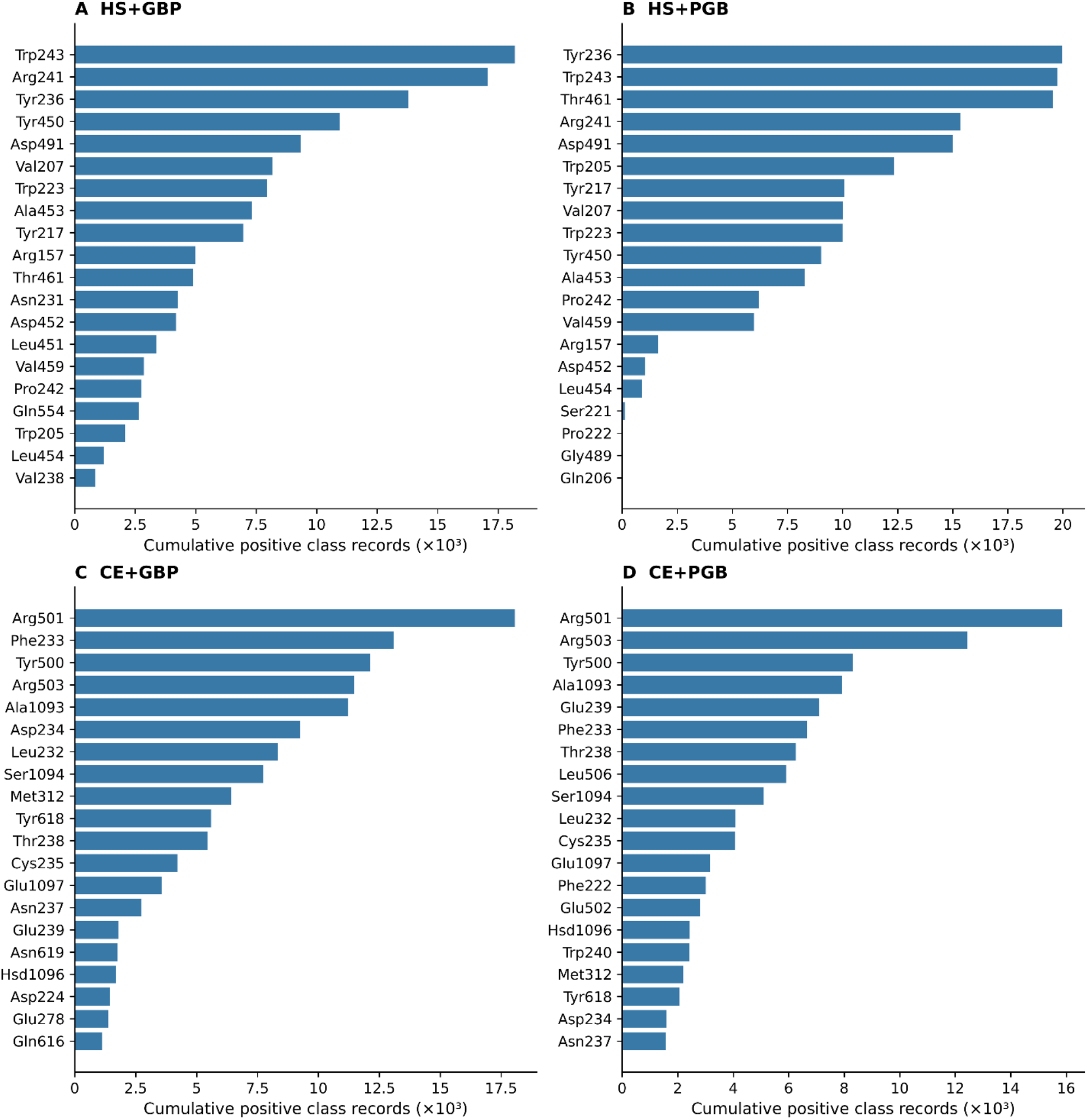
Cumulative positive contact-class records for the top 20 residues in each analyzed trajectory. Panels: (A) HS+GBP, (B) HS+PGB, (C) CE+GBP, and (D) CE+PGB. Counts sum binary class-presence records across all 5,001 frames and all contact classes, including generic proximity. A residue can contribute to several classes in one frame; counts may therefore exceed 5,001 and are not residue-occupancy percentages or numbers of independent bond-formation events. Residues are ordered by decreasing cumulative count. Complete counts for all recorded residues were retained with the analysis data.

Therefore, the rankings describe cumulative proximity classifications rather than residue occupancy or numbers of independently formed bonds. Arg501 had the largest cumulative class count in both UNC-36 trajectories. The next highest counts were Phe233, Tyr500, Arg503, and Ala1093 for CE+GBP, and Arg503, Tyr500, Ala1093, and Glu239 for CE+PGB. Phe233 and Tyr500 were recurrently close to the ligands, but their residue identities and distance-based classifications do not establish aromatic interaction geometry. Contacts involving Arg501 and Arg503 identify these residues as candidates for further investigation.

The selected-residue timelines in Figure 5 show changes in contact patterns for the ten residues with the highest cumulative counts in each trajectory. Some contact classes persisted over extended periods, while others appeared intermittently. Generic proximity accounted for the largest share of recorded contacts in all four systems: 40.4%, 38.7%, 40.5%, and 44.4% for HS+GBP, HS+PGB, CE+GBP, and CE+PGB, respectively (Figure 6). In HS+GBP, ligand N/H-to-protein O/N proximity accounted for 17.4% of records, followed by aromatic-residue proximity at 13.8% and ligand O/N-to-protein N/H proximity at 12.7%. In HS+PGB, aromatic-residue and O/N-to-N/H proximity accounted for 16.7% and 15.3%, respectively. The atom selections and distance cutoffs used for these classes are provided in Table S4.

**Figure 5.**
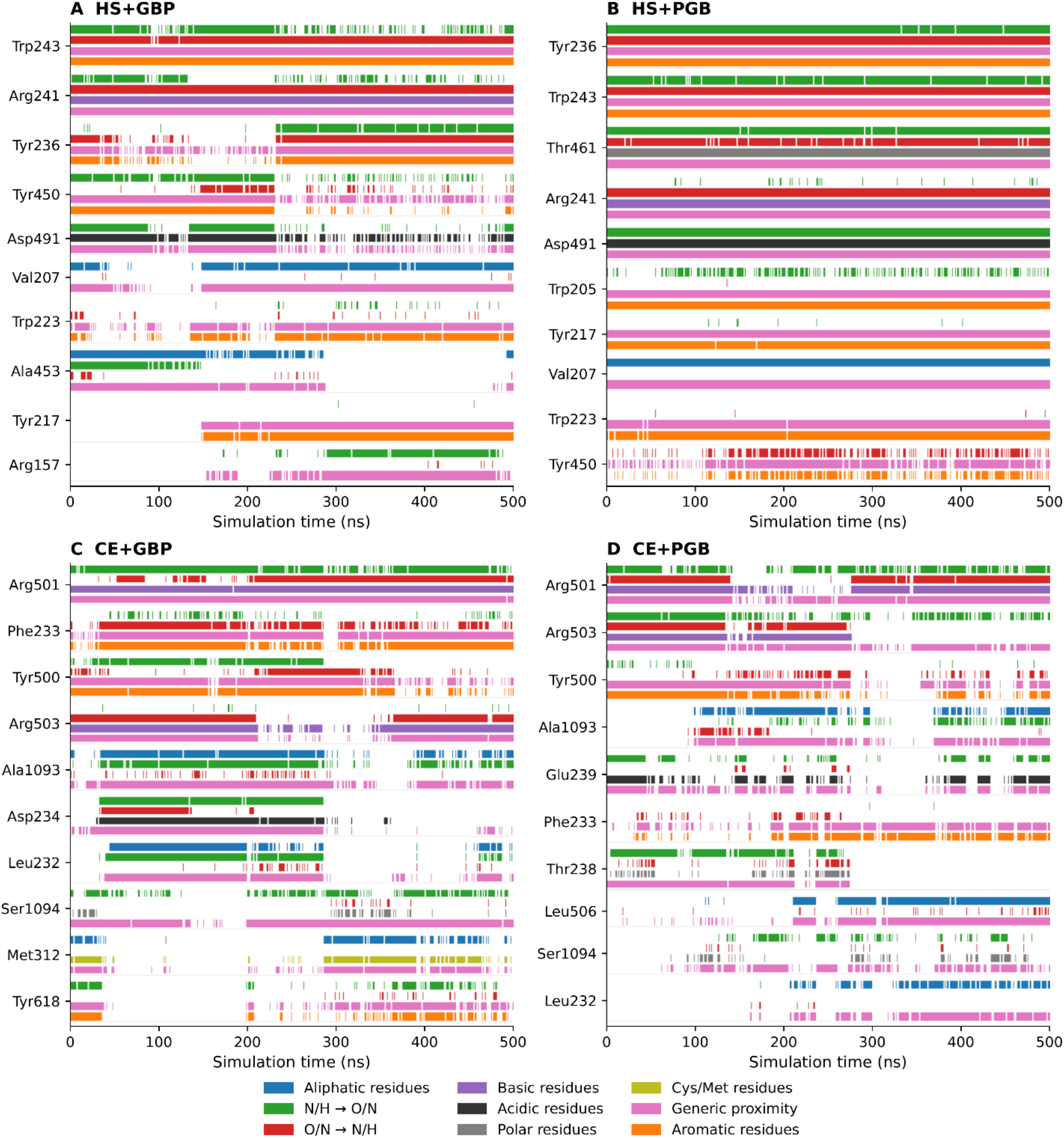
Contact-class timelines for the ten residues with the largest cumulative counts in each trajectory. Panels: (A) HS+GBP, (B) HS+PGB, (C) CE+GBP, and (D) CE+PGB. Each residue has separate colored subrows for its positive contact classes, defined in Table S4. Each sampled frame is displayed as a centered 0.1-ns bin, clipped at the trajectory endpoints; adjacent positive bins are joined for display. These segments describe presence at the saved sampling interval and do not establish chemical-bond lifetimes between frames. Complete timelines for all recorded residues were retained with the analysis outputs.

**Figure 6.**
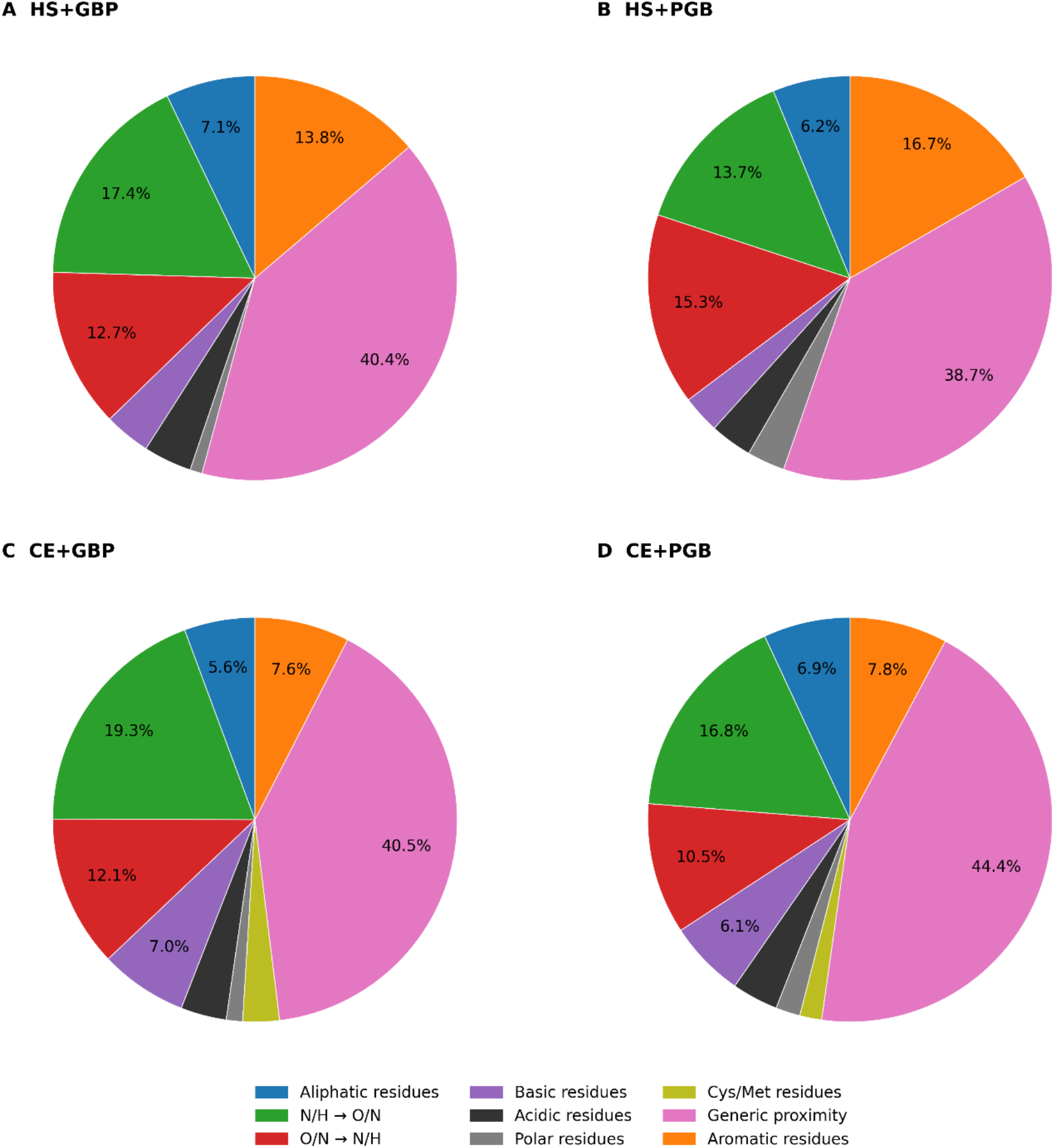
Distribution of recorded distance-based contact classes. Panels: (A) HS+GBP, (B) HS+PGB, (C) CE+GBP, and (D) CE+PGB. Percentages use all positive residue–class–frame records from all 5,001 frames, including residues omitted from the selected displays in Figures 3–5. Classes overlap and the percentages are not mutually exclusive physical interaction types or energetic contributions. N/H-to-O/N and O/N-to-N/H labels give the ligand atom-name selection first and the protein selection second. Labels below 5.0% are omitted from the chart for readability; their records remain included in the totals. Exact selections and cutoffs are listed in Table S4.

In CE+GBP, N/H-to-O/N and O/N-to-N/H proximity accounted for 19.3% and 12.1% of records, respectively, while basic-residue and aromatic-residue proximity accounted for 7.0% and 7.6%. In CE+PGB, N/H-to-O/N proximity represented 16.8%, basic-residue proximity 6.1%, and aromatic-residue proximity 7.8%. The UNC-36 trajectories therefore showed larger basic-residue proximity fractions than the human trajectories, where these fractions were 3.7% for HS+GBP and 3.0% for HS+PGB, and smaller aromatic-residue fractions. These percentages summarize positive contact-class records across residues and frames. Because a residue can contribute to more than one class in the same frame, they describe overlapping proximity classifications rather than distinct chemical interaction types.

### 2.5. PCA and free-energy landscapes describe the sampled conformational distributions

PCA showed that PC1 and PC2 together captured 45.4% of positional variance in HS+GBP and 43.1% in HS+PGB (Figure 7A,D), compared with 69.8% in CE+GBP and 61.4% in CE+PGB (Figure 7G,J). PC1 alone accounted for 58.4% of the variance in CE+GBP and 49.4% in CE+PGB, whereas the corresponding values were 32.2% and 29.4% in the two human systems. The leading collective motions were therefore concentrated into fewer dominant components in the UNC-36 trajectories. The PC1-PC2 projections and FELs also differed in topology (Figure 7B,C,E,F). HS+GBP sampled several partially separated conformational regions and multiple low-energy basins while HS+PGB followed a more elongated distribution with major regions connected by intermediate conformations. CE+GBP traced a narrower trajectory dominated by PC1, whereas CE+PGB showed a branched trajectory with a prominent transition along PC2 (Figure 7H,K). Together, these analyses show that conformational variance was less concentrated in the first two principal components in the human systems than in the UNC-36 systems. Consequently, the PC1– PC2 projections provide a less complete representation of the human trajectories than of the UNC-36 trajectories. CE+GBP sampling occurred predominantly along PC1, whereas CE+PGB showed a clearer transition along PC2. These distributions describe the motions sampled in individual trajectories and do not establish reproducible species- or ligand-dependent differences in stability or function.

**Figure 7.**
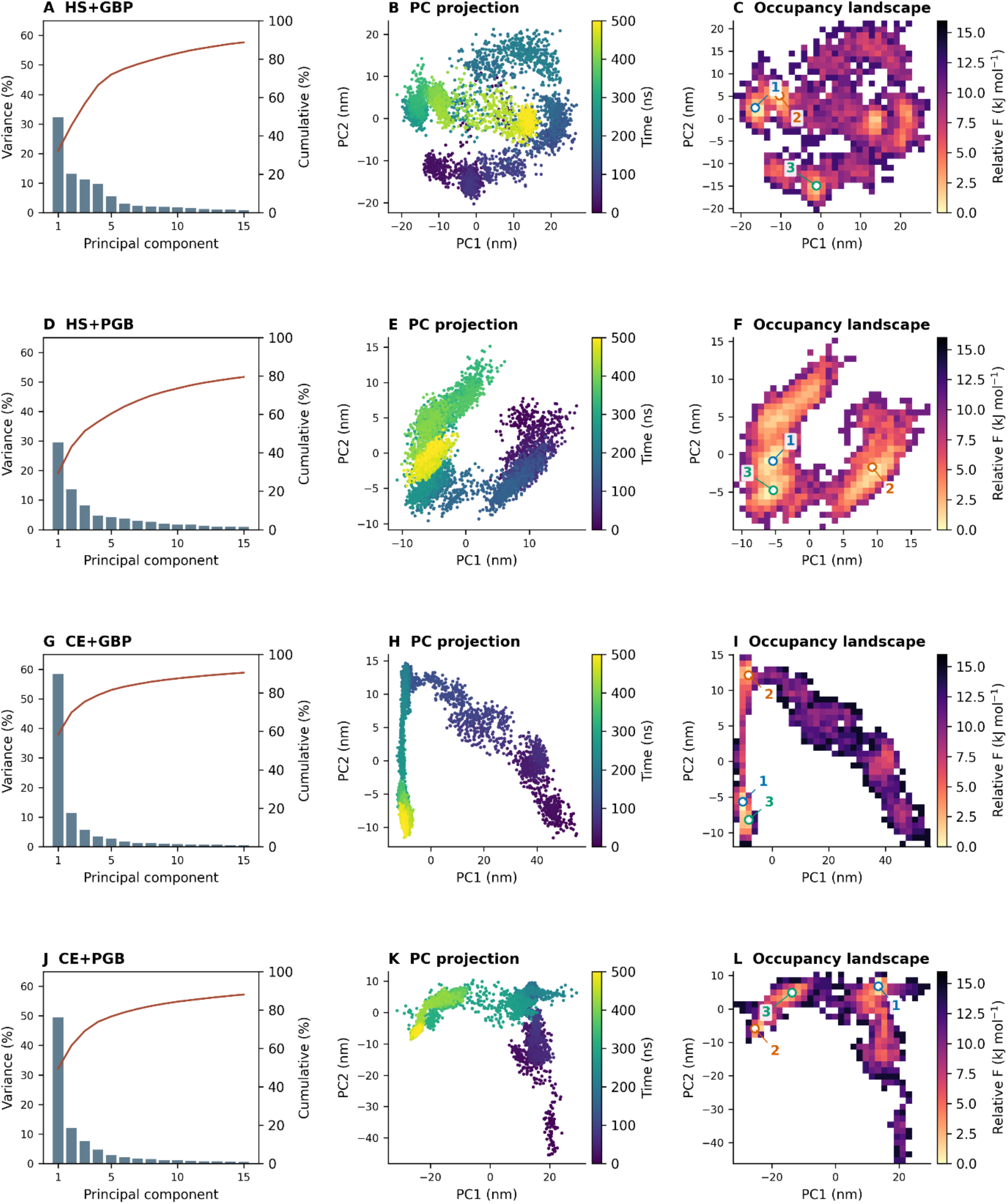
Principal component and occupancy-based free-energy landscape analyses of human α2δ-1 and *C. elegans* UNC-36 gabapentinoid complexes. (A, D, G, J) Variance explained by the first 15 principal components (bars, left axes) and cumulative variance (lines, right axes) for HS+GBP, HS+PGB, CE+GBP, and CE+PGB. (B, E, H, K) PC1–PC2 projections colored by simulation time. (C, F, I, L) Relative occupancy-derived energies on the original 32 × 32 grids, recalculated at 303.15 K; white bins were unoccupied. Numbered markers identify the actual frames selected from the three lowest-energy grid minima, ordered by increasing bin energy within each system. Their structures and pairwise comparisons are shown in Figure S2. PC1 and PC2 together accounted for 45.4%, 43.1%, 69.8%, and 61.4% of total variance in HS+GBP, HS+PGB, CE+GBP, and CE+PGB, respectively. Each system has its own PC basis; coordinates are not directly comparable between systems. These landscapes describe the sampled trajectories and do not establish convergence, equilibrium populations, or independent metastable states.

Representative structures were extracted from the three lowest-energy grid minima in each landscape (Figure 7C,F,I,L; Figure S2, Table S6). Pairwise whole-protein backbone RMSDs among these snapshots ranged from 2.84–5.00 Å for HS+GBP, 2.87–3.68 Å for HS+PGB, 3.01– 4.18 Å for CE+GBP, and 4.45–7.04 Å for CE+PGB. After local backbone alignment, ligand heavy-atom RMSDs ranged from 1.31–5.70, 0.53–0.76, 4.98–7.08, and 3.82–6.94 Å, respectively. Thus, the three selected HS+PGB snapshots retained similar local ligand poses despite differences across the full protein. HS+GBP minima 1 and 2 were more similar locally than either was to minimum 3, the earlier 44.3-ns snapshot. Both UNC-36 systems showed larger ligand rearrangements among the selected structures. These comparisons describe selected conformations within each trajectory; grid minima were not assumed to represent independent metastable states.

### 2.6. MM/GBSA estimates differ among the four analyzed trajectories

MM/GBSA calculations over the final 300-500 ns gave mean binding-energy estimates of −37.70 ± 3.08 kcal/mol for HS+PGB, −24.76 ± 3.42 kcal/mol for HS+GBP, −22.00 ± 8.50 kcal/mol for CE+GBP, and −12.08 ± 6.04 kcal/mol for CE+PGB (mean ± sample SD across 501 frames; Figure 8). These estimates exclude an explicit conformational entropy contribution, and the reported SD describes variation among the sampled frame values. CE+GBP showed substantial temporal variation, with mean estimates of −12.03, −24.86, −27.39, and −23.67 kcal/mol across four consecutive 50-ns intervals (Table S3). Time-window and block analyses were performed for all four systems (Table S3; Figure S1). CE+GBP showed substantial variation over time, making its overall mean less certain.

**Figure 8.**
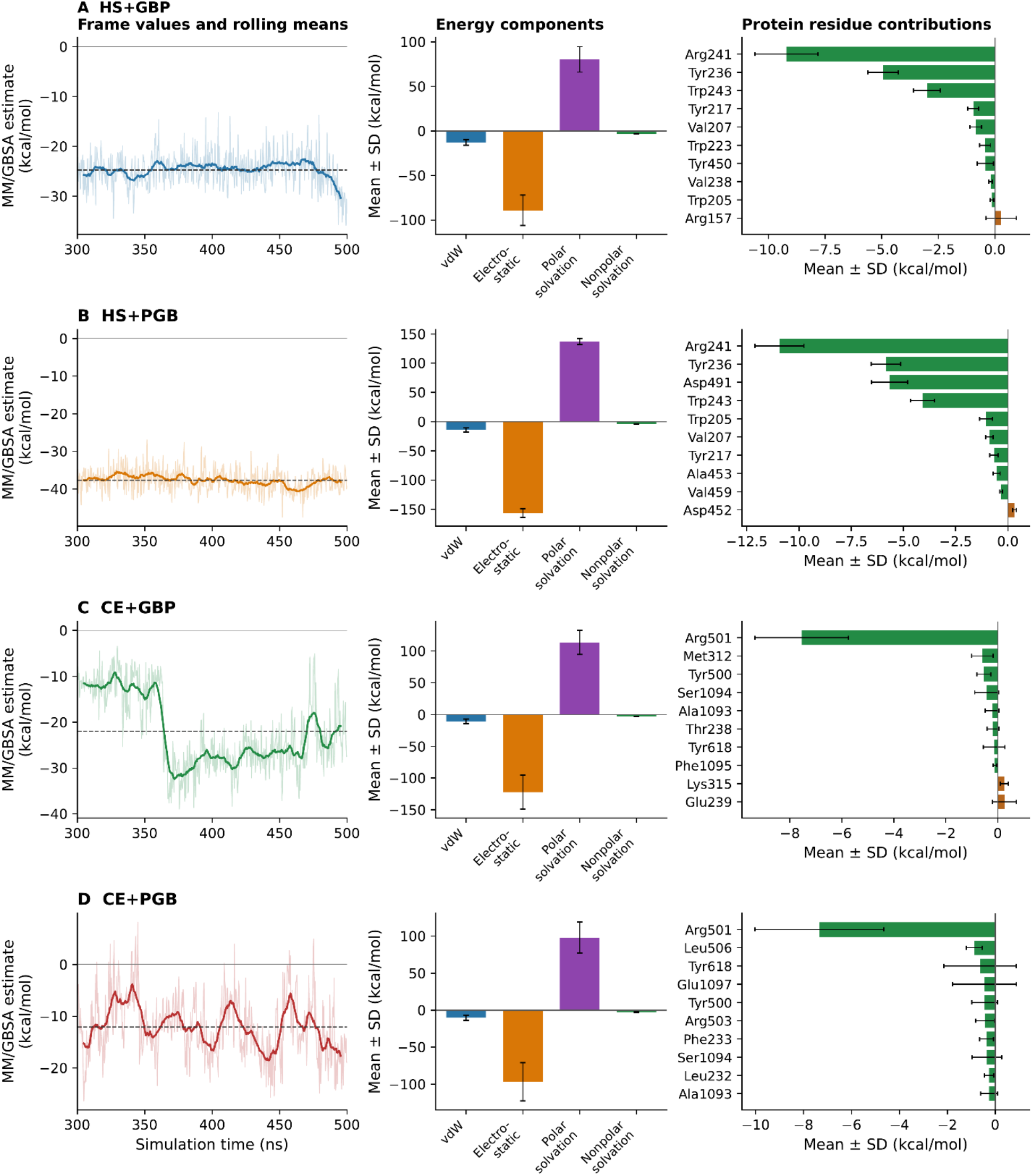
MM/GBSA energy estimates and residue decomposition for 300–500 ns, without an explicit conformational entropy contribution. Rows: (A) HS+GBP, (B) HS+PGB, (C) CE+GBP, and (D) CE+PGB. Left: 501 frame values (pale lines), centered 25-frame rolling means (colored lines), and full-window means (dashed lines). Center: mean van der Waals, electrostatic, polar-solvation, and nonpolar-solvation contributions. Right: the ten protein residues with the largest absolute mean contributions among the saved total-decomposition selections, ordered by signed mean. All component and residue error bars show sample SD across 501 paired-frame complex-minus-receptor-minus-ligand values. SD describes the spread of sampled values, not uncertainty in a mean, a confidence interval, or variability across independent replicas.

Within the analyzed trajectories, per-residue interaction energies were calculated for the selected pocket residues (Figure 8). Arg241 was among the strongest favorable contributors in the human complexes, whereas Arg501 was the dominant favorable protein contributor in both UNC-36 systems.

## 3. Discussion

Structural comparison identified close local backbone correspondence within a selected pocket core of human α2δ-1 and UNC-36. The modeled human pocket agreed with the experimental rabbit α2δ-1 gabapentin-binding region (PDB 8FD7), with a Cα RMSD of 0.63 Å across 14 mapped residues. For the human and UNC-36 models, the Cα RMSD was 9.07 Å across the complete 14-residue mapping and 0.53 Å across the selected 13-residue core after excluding Thr461/Ala513. The approximately 36.3 Å separation of this pair after core alignment limits extrapolation from the selected core to the complete mapped binding region. The low pLDDT of UNC-36 Ala513 (44.25) and elevated PAE involving this position indicate uncertainty in its predicted structure and relative placement. Its displacement therefore cannot, by itself, establish a biological structural difference. These structural comparisons provide a basis for examining how differences in pocket residues relate to the ligand-contact patterns observed in the simulations.

This interpretation is consistent with the established pharmacology and structural biology of mammalian α2δ-1. Gabapentin was first identified as a high-affinity α2δ ligand (6), and mutagenesis subsequently established a conserved arginine as critical for binding (12); historically designated Arg217, it corresponds to Arg241 in the full-length human CACNA2D1 sequence used here, as the earlier numbering excludes the 24-residue N-terminal signal peptide. The first dCache_1 domain contains a conserved amino acid recognition site (8), and cryo-EM later resolved GBP fully enclosed within this pocket (8), with mirogabalin-bound human α2δ-1 confirming the region as a common gabapentinoid recognition site (29). In the analyzed human trajectories, Arg241, Tyr236, Trp243, Tyr450, Asp491, and adjacent residues contacted GBP or PGB repeatedly within the modeled mammalian binding region. Within the analyzed human trajectories, PGB showed lower ligand RMSD, higher contact counts, and a more negative MM/GBSA estimate than GBP. PGB is experimentally characterized as a potent and selective ligand of α2δ-1 and α2δ-2 (7). In the analyzed UNC-36 trajectories, the modeled contact pattern differed from that in the human trajectories despite local backbone correspondence within the selected core. Arg501 had the largest cumulative proximity-class count in both CE+GBP and CE+PGB, with Arg503 also contributing recurrent contacts. Tyr500 and Phe233 were frequently close to the ligands. Compared with the human trajectories, the UNC-36 trajectories showed a larger proportion of basic-residue proximity records and a smaller proportion of aromatic-residue proximity records. The sequence and structural comparisons offer a possible explanation for this pattern. Human Leu451 and Ala453 correspond to Arg501 and Arg503 in UNC-36, whereas human Arg241 corresponds to Thr238. These substitutions redistribute positively charged residues within the corresponding pocket region. Several aromatic positions are also identical or conservatively substituted, including Tyr217/Tyr215, Trp223/Trp221, Tyr236/Phe233, Trp243/Trp240, Tyr450/Tyr500, and Trp205/Phe203. Together, these features suggest that similarities in the selected pocket backbone coexist with differences in local charge distribution that may influence ligand contacts. Comparable sensitivity to changes in pocket chemistry has been described among mammalian α2δ isoforms, where side-chain volume, polarity, and hydrogen-bonding capacity influence gabapentinoid recognition (11). Our simulations therefore suggest a more arginine-centred contact pattern in UNC-36, with Arg501 and Arg503 emerging as candidates for experimental investigation. This molecular interpretation provides a useful extension of our previous behavioral findings. We previously showed that mutations in *unc-2* and *unc-36* impaired thermal avoidance to noxious heat and that GBP and PGB reduced thermal nocifensive behavior in *C. elegans*. These behavioral effects were accompanied by proteomic changes in pathways associated with VGCC function and nociceptive processing (18). Those findings linked a functional relationship between gabapentinoids, VGCC-associated pathways, and thermal nociception, but did not establish direct binding to UNC-36. The present study contributes a molecular hypothesis by identifying a candidate pocket in UNC-36 with a selected backbone core corresponding to human α2δ-1 and modeled contacts with both ligands. Arg501, Arg503, Tyr500, and Phe233 showed the highest contact frequencies, identifying them as priority targets for experimental validation.

A similar relationship between conserved structure and different ligand-contact chemistry has emerged from previous nociception studies in our laboratory. Capsaicin and related vanilloids reduce thermal nocifensive responses in *C. elegans*, with the TRPV-like proteins OCR-2 and OSM-9 contributing to these responses in a compound-dependent manner (21–23). More recently, comparative MD analysis showed that mammalian TRPV1 and *C. elegans* OSM-9/OCR-2 preserve common features of the capsaicin-recognition architecture while using different residue-level interactions to engage the ligand (25). These findings support the use of *C. elegans* to investigate potentially shared pharmacological mechanisms, while indicating that species-specific features of the molecular target may limit the transfer of compound-specific conclusions, such as potency or selectivity, to mammalian systems.

The conformational and energetic analyses further distinguished the four complexes. PC1 and PC2 together accounted for 45.4% and 43.1% of the positional variance in HS+GBP and HS+PGB, respectively, compared with 69.8% in CE+GBP and 61.4% in CE+PGB (Figure 7). The higher contribution of the first two components in the UNC-36 systems indicates that a larger proportion of their sampled motion was concentrated into a few dominant collective modes (30). This was particularly evident for CE+GBP, in which PC1 alone accounted for 58.4% of the variance and the trajectory followed a more directional distribution in PC1–PC2 space. In contrast, HS+GBP sampled several partially separated conformational regions, HS+PGB showed a more elongated distribution, and CE+PGB displayed a broader, branched pattern of conformational sampling. The corresponding FELs reflected these differences in the regions sampled during the trajectories, although the PCA and FEL analyses were not used to assign specific functional states. Comparison of structures associated with the FEL minima helped distinguish changes in the overall protein conformation from changes in ligand positioning. The selected HS+PGB snapshots retained similar local ligand poses, whereas the other systems showed larger differences after local protein alignment (Table S6; Figure S2). MM/GBSA analysis over the final 200 ns gave binding-energy estimates of −37.70 ± 3.08 kcal/mol for HS+PGB, −24.76 ± 3.42 kcal/mol for HS+GBP, −22.00 ± 8.50 kcal/mol for CE+GBP, and −12.08 ± 6.04 kcal/mol for CE+PGB (mean ± sample SD across frames). The SDs describe variation among the sampled values. Within this comparison, PGB had a more favorable mean estimate than GBP in human α2δ-1, whereas the opposite pattern was observed in UNC-36. Together, the structural and energetic analyses suggest differences in how the two ligands are accommodated by the modeled proteins.

For future ligand design, our findings suggest testing related gabapentinoid analogues that retain the amine and carboxylate groups while varying their spacing, stereochemistry, and hydrophobic substituents. The human Leu451/UNC-36 Arg501 and human Ala453/UNC-36 Arg503 differences suggest that local charge distribution could influence ligand recognition, a possibility that could be explored through cross-species structure–activity comparisons. Candidate compounds could be evaluated using direct binding or target-engagement measurements, together with VGCC functional assays. In UNC-36, individual and combined Arg501→Leu and Arg503→Ala substitutions could help determine the contribution of these residues to ligand recognition, with the reciprocal human substitutions providing a complementary test. These experiments should include controls for receptor expression, localization, and basal channel function. Thermal-avoidance assays, accompanied by locomotor controls, could then connect the molecular findings to whole-animal responses and help distinguish changes in drug recognition from broader effects on sensory or motor function.

Together, these findings have implications for the use of *C. elegans* in early-stage drug screening. Whole-organism chemical screening studies in *C. elegans* can identify bioactive compounds while retaining information about their effects on conserved physiological pathways. For instance, Nemadipine-A, a widely used calcium blocker, illustrates a similar cross-species relationship - a screening of 14,100 small molecules in *C. elegans* identified nemadipine-A based on marked morphological and egg-laying defects, and a subsequent suppressor screening identified *egl-19*, the sole L-type calcium channel α1-subunit gene in *C. elegans*, as its target. Importantly, Nemadipine-A was later shown to antagonize vertebrate L-type calcium channels, supporting conservation of the pharmacological target across species (31). This example is relevant to the present study because it shows that *C. elegans* VGCC components can retain pharmacologically meaningful relationships with their vertebrate counterparts, supporting the broader use of *C. elegans* in high-throughput screening for bioactive compounds and ion-channel pharmacology (14,32). Our previous thermal-avoidance work provides a behavioral readout for nociceptive pharmacology (18), while the present study identifies candidate residues involved in gabapentinoid recognition. Structural similarities within the selected pocket core provide a rationale for investigating UNC-36 as a potential gabapentinoid target. Differences in residue identity and the uncertain placement of Ala513 highlight features to examine when comparing ligand recognition in UNC-36 and human α2δ-1. A compound that modifies nocifensive behavior in *C. elegans* should not automatically be expected to interact with human α2δ-1 through the same residues or with the same affinity. A practical strategy would therefore use *C. elegans* for initial whole-organism screening of compounds acting on conserved nociceptive pathways, followed by validation of binding and selectivity at the human target.

Several limitations should be considered when interpreting these findings. The receptor structures were predicted, and uncertainty around UNC-36 Ala513 limits interpretation of its displacement relative to human Thr461. The close backbone correspondence applies to the selected 13-residue core rather than the complete mapped binding region. One final 500-ns trajectory was analyzed per complex, with starting poses selected using docking scores, contact inspection and, in some cases, preliminary MD observations. The comparisons therefore describe behavior from those selected configurations; independent replicas and alternative starting poses would be needed to assess reproducibility.

The MM/GBSA estimates exclude an explicit conformational entropy contribution and remain sensitive to sampling and model assumptions, particularly given the temporal variation observed for CE+GBP. Reported SDs describe variation across frames rather than uncertainty in the means, and the estimates do not establish experimental affinity differences. The custom proximity profiles describe distance-based contacts rather than chemically specific interactions. Direct binding or target-engagement measurements, supported by controlled mutagenesis and functional assays, are needed to test the proposed role of UNC-36 and its candidate residues in gabapentinoid recognition and behavioral responses.

## 4. Conclusion

The modeled human α2δ-1 and UNC-36 structures show close Cα correspondence within a selected 13-residue core, while the complete 14-residue mapping includes an uncertain, displaced Thr461/Ala513 pair. The 0.53 Å core RMSD therefore describes local backbone correspondence; the full mapping yielded 9.07 Å. One final 500-ns trajectory per complex revealed differences in recorded ligand contacts, sampled conformations, and MM/GBSA energy scores. These observations identify Arg501 and Arg503 and the surrounding modeled environment as candidates for tests of gabapentinoid recognition by UNC-36. They provide hypotheses for analogue design and comparative pharmacology, pending direct binding and functional validation.

## 5 Materials and methods

### 5.1. Protein sequences, structural models, and cross-species comparison

The human voltage-gated calcium-channel α2δ-1 sequence (CACNA2D1; UniProt P54289) and the *C. elegans* UNC-36 sequence (UniProt P34374) were retrieved from the UniProt database. Then, three-dimensional structures were independently predicted from their respective amino-acid sequences using ColabFold v1.5.5, which implements AlphaFold2 for protein structure prediction (33,34). Five ColabFold models per protein were evaluated using MolProbity. The model with the highest Ramachandran-favored fraction was selected for each protein. The selected structures were refined using the GalaxyRefine server (35). Human model 3 and UNC-36 model 4 were used for subsequent receptor preparation. Model quality was then evaluated using complementary structure-validation tools. Overall model quality was assessed using ERRAT and ProSA-web, while stereochemical quality was examined by Ramachandran plot analysis and MolProbity (Table S2). Local prediction confidence was assessed using Predicted local distance difference test (pLDDT) and predicted aligned error (PAE) values from the original ColabFold outputs for the complete 14-residue mapping and the selected 13-residue core. Mean PAE included both directions for each residue pair and excluded self-pairs. The core was selected based on structural correspondence rather than confidence scores. These values describe the original predictions, whereas the post-refinement geometry assessments are reported separately (Tables S1 and S2).

The amino-acid sequences of human α2δ-1 and *C. elegans* UNC-36 were initially compared using Clustal Omega to verify residue correspondence within the gabapentinoid-binding region and to support mapping of experimentally defined mammalian α2δ-1 pocket residues onto UNC-36. Following the MD simulations, a refined sequence comparison was performed with MAFFT version 7 the L-INS-i strategy (36). Residues that formed recurrent contacts with GBP or PGB during the final trajectories were mapped onto the MAFFT alignment to assess conservation of the ligand-interacting environment. Positions were classified as identical when the same amino acid occurred at aligned positions and as conservative substitutions when residues retained similar physicochemical properties. This analysis was used to distinguish strict residue conservation from broader conservation of the gabapentinoid-binding environment. Moreover, structural alignment was performed in PyMOL to compare the human α2δ-1 and UNC-36 extracellular ligand-binding regions. Whole-protein overlays were used for orientation, whereas interpretation focused on the local region corresponding to the mammalian dCache1 gabapentinoid-binding site defined structurally by Chen et al. (2023). Because automated domain annotations differ between human α2δ-1 and UNC-36, the UNC-36 site is described here as a structurally corresponding or dCache1-like region rather than as an experimentally validated gabapentinoid-binding domain.

The experimentally resolved rabbit CaVα2δ-1–gabapentin complex (PDB 8FD7) was used as a structural reference for the gabapentinoid-binding site (11). Fourteen α2δ-1 residues located within 5 Å of gabapentin were identified in PyMOL and mapped onto human α2δ-1 (CACNA2D1) and subsequently onto UNC-36 according to the sequence alignment. Local structural conservation was evaluated by pairwise superposition of corresponding Cα atoms in PyMOL. The modeled human pocket was first compared with the experimental rabbit CaVα2δ-1–gabapentin complex (8FD7) structure as an internal structural validation. Human and UNC-36 pocket residues were then compared using the same residue mapping. Because the sequence-aligned human Thr461/UNC-36 Ala513 pair occupied different spatial positions, RMSD was calculated for both the complete 14-residue mapping and the selected 13-residue pocket core. The core RMSD describes local backbone similarity rather than conservation of the entire binding region or side-chain geometry.

### 5.2 Ligand preparation and molecular docking

Three-dimensional structures of GBP and PGB were obtained from PubChem and prepared for docking. Flexible docking was performed with the AutoDockFR workflow, which allows explicitly specified receptor-side-chain flexibility (37). The human docking region was guided by the mammalian α2δ-1 gabapentin-binding site identified by mutagenesis and cryo-EM studies (11,12). The corresponding UNC-36 region was defined by sequence mapping and structural superposition. Each search included 100 genetic-algorithm evolutions, with a maximum of 3,000,000 evaluations per evolution, and the resulting solutions were clustered using a 2.0 Å RMSD cutoff. Starting poses were chosen based on docking scores, visual inspection of pocket occupancy, ligand orientation and protein–ligand contacts, and, in some cases, preliminary MD observations. The selected poses and their docking scores are reported in Table S5.

### 5.3 Molecular-dynamics system preparation and simulations

Molecular dynamics (MD) simulations were performed with GROMACS 2024.4 on the high-performance computing resources provided by the Digital Research Alliance of Canada (38). The protein–ligand complexes were prepared using the Solution Builder module of CHARMM-GUI and parameterized with the CHARMM36 force field (39,40). Each system was solvated with TIP3P water, neutralized, and adjusted to an ionic strength of 0.150 M KCl. After system construction, energy minimization was carried out using the steepest-descent algorithm, followed by the multistep equilibration protocol generated by CHARMM-GUI with progressively reduced positional restraints. MD production simulations were run under periodic boundary conditions using the Verlet cutoff scheme and a 2-fs integration time step. Short-range electrostatic interactions were computed with a 1.2 nm cutoff, and long-range electrostatics were treated with the particle-mesh Ewald (PME) method. Van der Waals interactions were force-switched between 1.0 and 1.2 nm, with a neighbor-list cutoff of 1.2 nm. Temperature was maintained at 303.15 K with the velocity-rescale thermostat (τ = 1.0 ps), and pressure was maintained at 1 bar with the C-rescale barostat using isotropic coupling (τ = 5.0 ps; compressibility = 4.5 × 10⁻⁵ bar⁻¹). Bonds involving hydrogen atoms were constrained using the LINCS algorithm, which permitted a 2-fs time step. Each final production trajectory was initiated with a 1-ns run and extended from checkpoint files to a cumulative duration of 500 ns. These extensions continued the same trajectory and did not constitute independent replicas. Compressed trajectory coordinates were written every 100 ps for subsequent analyses. Ligand parameters were assigned using the CHARMM General Force Field (CGenFF), with both ligands represented in their zwitterionic forms with zero net charge.

### 5.4 Molecular dynamics trajectory analysis

Before analysis, trajectories were corrected for periodic boundary effects, centered on the protein, and aligned to remove overall rotation and translation. Analyses covered the full 500-ns trajectories. Protein backbone RMSD was calculated after backbone alignment using the production starting structure as reference. Ligand heavy-atom RMSD was calculated relative to the first frame of the protein-aligned trajectory, without additional ligand fitting. Protein radius of gyration and whole-protein–ligand centre-of-mass distances were calculated using GROMACS. Protein–ligand hydrogen bonds were counted using gmx hbond-legacy, with a donor–acceptor distance cutoff of 0.35 nm and a hydrogen–donor–acceptor angular cutoff of 30° (0° corresponding to an extended arrangement). Protein–ligand contacts were counted as atom pairs within 0.35 nm, including hydrogen atoms.

PCA was performed using the GROMACS tools gmx covar and gmx anaeig, with backbone atoms (N, Cα and C) used for least-squares fitting and covariance analysis without mass weighting. All 5,001 frames from 0–500 ns, saved at 100-ps intervals, were included. Each system was analyzed separately, and trajectories were projected onto their first two principal components.

FELs were calculated from frame occupancies on 32 × 32 PC1–PC2 grids using Fᵢ = −RT ln(nᵢ/nₘₐₓ), where nᵢ is the number of frames in a bin, nₘₐₓ is the highest bin count, R is the molar gas constant, and T = 303.15 K. Empty bins were excluded, and no smoothing was applied.

Representative structures were selected from the three lowest-energy grid minima in each system. For each minimum, the trajectory frame within that bin nearest its centre was selected using PC coordinates normalized by bin width. Pairwise backbone RMSDs were calculated after whole-protein and local backbone alignment. For local alignment, a common residue set was used across the three snapshots of each system, comprising protein residues with a heavy atom within 8 Å of a ligand heavy atom in any selected snapshot. Ligand heavy-atom RMSDs were calculated after local protein alignment, matching atoms by name without additional ligand fitting or symmetry correction. Snapshot times and structural overlays are presented in Figure S2.

Protein-ligand proximity contacts were profiled using custom Python scripts with MDAnalysis (41,42) and SciPy pairwise distance calculations. The same contact rules were used for all four systems (Table S4). Each centered trajectory contained 5,001 analyzed frames spanning 0–500 ns. Candidate protein residues were identified by a 7 Å proximity screen applied every fifth frame; the union of these residues was then evaluated across all frames. For each residue, contact class, and frame, a binary presence value was assigned when any selected ligand–protein atom pair was closer than the class-specific cutoff (3.5–5.0 Å); otherwise the value was zero. Atom-name and residue selections are listed in Table S4. For residue-based protein selections, all atoms of those residues were eligible, including backbone atoms and hydrogens when present. These classes represent proximity rather than chemically specific interactions. The Ser/Thr and Asn/Gln subclasses were combined into one polar-residue category for plotting. For each residue and class, contact frequency was calculated as the percentage of analyzed frames with a presence value of one. Cumulative residue counts summed these binary values across frames and classes; a residue could therefore contribute to multiple classes in the same frame. Heatmaps display the number of positive classes for each residue and frame, while timelines show the frames in which each class was present. Category percentages were calculated from all positive residue–class–frame records and describe the distribution of overlapping proximity classifications. Figure 3 includes up to 20 residues with at least one nongeneric record, Figure 4 shows the top 20 recorded residues, and Figure 5 shows the top ten. Residues were ranked by cumulative counts across all classes within each trajectory. Figure 6 includes all positive records without this display selection.

### 5.5 MM/GBSA energy estimates, residue decomposition, and sampling diagnostics

Molecular Mechanics/Generalized Born Surface Area (MM/GBSA) calculations were performed using gmx_MMPBSA v1.5.0.3 (43,44). For each system, 501 evenly spaced frames from 300–500 ns were analyzed at 400-ps intervals. Receptor and ligand coordinates were extracted from the same complex frames, and their topologies were derived from the supplied CHARMM complex topology, using the single-trajectory end-point protocol. The reported energy score was calculated as:

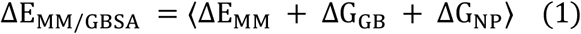

Here, Δ denotes the difference between the complex and the separate receptor and ligand for each energy term. E_MM_ is the molecular-mechanics energy, G_GB_ is the polar solvation contribution, and G_NP_ is the nonpolar solvation contribution. Angle brackets denote the arithmetic mean over the analyzed frames. No explicit conformational entropy contribution was included. The nonpolar solvation energy for each molecular species was modeled as:

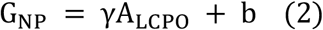

Here, A_LCPO_ is the solvent-accessible surface area estimated using the LCPO method, γ is the surface-area coefficient, and b is the constant offset. The surface-area coefficient γ was 0.0072 kcal/mol Å⁻², and the offset b was 0 kcal/mol. Polar solvation calculations used the OBC-II model (igb = 5), mbondi2 atomic radii and a salt concentration of 0.150 M. Internal and external dielectric constants were 1.0 and 78.5, respectively. Unspecified settings retained the gmx_MMPBSA v1.5.0.3 defaults.

Related computational studies have used MD, free-energy perturbation and MM/GBSA to investigate ligand recognition by guanine riboswitch mutants (45), and Gaussian-accelerated MD to examine ligand recognition by an SAM-III riboswitch (46). For the present MM/GBSA calculations, the OBC-II generalized Born model (igb = 5) was used as an established, computationally efficient approximation for estimating polar solvation contributions (47). The same model and settings were applied to all four systems to maintain a consistent calculation protocol. Sensitivity to alternative GB models or dielectric settings was not evaluated. MM/GBSA binding free-energy estimates were calculated without an explicit conformational entropy contribution.

MM/GBSA binding-energy estimates were reported as the mean ± sample SD across 501 frames from 300–500 ns. SD describes variation among frame values rather than uncertainty in the mean. Temporal correlation and time dependence were examined using autocorrelation, 50-ns averages and block analysis, with details provided in Table S3 and Figure S1.

Per-residue decomposition used idecomp = 2 and a 5-Å residue selection defined from the setup structure. Figure 8 shows the ten residues with the largest absolute mean contributions, with error bars representing sample SD across the analyzed frames.

### 5.6 Visualization and reporting

Protein structures and binding-site views were prepared in PyMOL. Interaction plots, residue-frequency summaries, MM/GBSA comparisons, PCA projections, and FEL panels were generated with reproducible Python workflows. The same labeling conventions were used across all four systems. For trajectory analyses, we used the complete 500-ns production simulations, whereas MM/GBSA analyses used the final 200 ns.

## Acknowledgements

This research was enabled in part by the High-Performance Computing infrastructure and support available at the Digital Research Alliance of Canada and Calcul Québec.

## Author’s contributions

JS, JDC and FB conceived and designed the research. JS, LM and JDC performed the experiments and analyzed the data. JS and JDC curated the data and wrote the original draft. FB and JdelC supervised the project. FB acquired funding and provided resources. All authors reviewed and approved the final manuscript.

## Data Availability

The data generated and analyzed during this study are available from the corresponding author upon reasonable request.

## Declaration of Generative AI Use

The authors used Claude (Anthropic) to assist with English-language editing and improve manuscript fluency. OpenAI Codex tools were used to assist in the development of code for verifying and re-analyzing existing molecular dynamics simulation outputs. All AI-generated outputs were reviewed, verified, and edited by the authors. The authors take full responsibility for the accuracy, integrity, and final content of the manuscript.

