## Supplemental data for "Structural conservation of the gabapentinoid binding site in human and *Caenorhabditis elegans* α2δ subunits: a docking and molecular dynamics perspective"

### Supplementary data

### Table S1: Pocket residue mapping and prediction confidence

#### A. Complete residue mapping and original model pLDDT

| Rabbit 8FD7 | Human | UNC-36 | Human pLDDT | UNC-36 pLDDT | Relationship |
|---|---|---|---|---|---|
| W207 | W205 | F203 | 93.94 | 95.81 | aromatic substitution |
| V209 | V207 | L205 | 95.88 | 97.75 | hydrophobic substitution |
| Y219 | Y217 | Y215 | 95.06 | 97.50 | identical |
| W225 | W223 | W221 | 79.50 | 89.12 | identical |
| Y238 | Y236 | F233 | 83.12 | 88.75 | aromatic substitution |
| R243 | R241 | T238 | 88.19 | 91.81 | loss of positive charge |
| W245 | W243 | W240 | 92.38 | 93.88 | identical |
| Y452 | Y450 | Y500 | 87.81 | 90.25 | identical |
| L453 | L451 | R501 | 84.44 | 86.56 | neutral → basic |
| D454 | D452 | E502 | 85.31 | 87.50 | acidic |
| A455 | A453 | R503 | 79.25 | 86.06 | neutral → basic |
| L456 | L454 | L504 | 79.44 | 87.12 | identical |
| T463 | T461 | A513 | 94.06 | 44.25 | Displaced pair; excluded from core |
| D493 | D491 | N568 | 93.94 | 96.00 | acidic → polar |

Rabbit UniProt P13806; human P54289; UNC-36 P34374. The sequence-mapped Thr461/Ala513 pair was excluded from the selected 13-residue core because it did not occupy corresponding positions after structural alignment.

#### B. Confidence summaries for the complete mapping and selected core

| Protein | Residue set | | Mean pLDDT | pLDDT range | Mean PAE (Å) |
|---|---|---|---|---|---|
| | Complete mapping | Selected core | | | |
| Human | 14 | | 88.02 | 79.25–95.88 | 2.34 |
| Human | | 13 | 87.56 | 79.25–95.88 | 2.37 |
| UNC-36 | 14 | | 88.03 | 44.25–97.75 | 3.75 |
| UNC-36 | | 13 | 91.39 | 86.06–97.75 | 1.99 |

pLDDT and PAE values were obtained from the original ColabFold predictions before refinement. Mean PAE includes both directions for each residue pair and excludes self-pairs. The 13-residue core was selected based on structural correspondence

**Table S2 Quality assessments of the refined receptor models**

| Protein | PROCHECK most favoured (%) | MolProbity Ramachandran-favoured (%) | ERRAT quality factor | MolProbity favoured rotamers (%) | ProSA Z-score |
|---|---|---|---|---|---|
| Human CACNA2D1 | 95.5 | 98.73 | 92.062 | 98.04 | −12.37 |
| *C. elegans* UNC-36 | 94.9 | 98.72 | 90.273 | 98.93 | −12.70 |

**Table S3 MM/GBSA time-window and block-size sensitivity**

### A. Mean estimates in consecutive 50 ns intervals

| System | 300–350 ns | 350–400 ns | 400–450 ns | 450–500 ns |
|---|---|---|---|---|
| HS+GBP | -25.33 | -24.09 | -24.33 | -25.23 |
| HS+PGB | -36.95 | -37.12 | -37.95 | -38.75 |
| CE+GBP | -12.03 | -24.86 | -27.39 | -23.67 |
| CE+PGB | -9.90 | -12.28 | -13.37 | -12.72 |

Values are in kcal/mol. Intervals include the left endpoint and exclude the right endpoint; each contains 125 frames at 0.4 ns spacing. The 500 ns endpoint is excluded only for equal-length blocking; full-window summaries use all 501 frames.

### B. Standard-error diagnostics from nonoverlapping block means (kcal/mol)

| System | 10 ns blocks<br>20 blocks | 20 ns blocks<br>10 blocks | 50 ns blocks<br>4 blocks |
|---|---|---|---|
| HS+GBP | 0.34 | 0.44 | 0.31 |
| HS+PGB | 0.26 | 0.32 | 0.41 |
| CE+GBP | 1.63 | 2.24 | 3.41 |
| CE+PGB | 0.75 | 0.82 | 0.76 |

Each value is the sample SD of block means divided by the square root of the number of blocks. These are sensitivity diagnostics, not established confidence intervals. The increasing values for CE+GBP do not establish a stable uncertainty estimate. Four 50 ns blocks provide limited information about longer-timescale variability. No explicit conformational entropy correction was included. See Figure S1.

**Table S4  Distance based contact rules**

The same rules were used for all four systems. Distances are in Å, and every cutoff uses a strict less-than comparison. Protein residue selections include all atoms of the listed residues.

| Contact class | Ligand selection | Protein selection | Cutoff (Å) |
|---|---|---|---|
| Aliphatic residues | name C* | ALA, VAL, LEU, ILE, MET, PRO | 4.0 |
| Aromatic residues | name C* | PHE, TYR, TRP | 4.5 |
| N/H to O/N proximity | name N* H* | name O* N* | 3.5 |
| O/N to N/H proximity | name O* N* | name N* H* | 3.5 |
| Ser/Thr residues | name O* N* | SER, THR | 4.0 |
| Asn/Gln residues | name O* N* | ASN, GLN | 4.0 |
| Acidic residues | name N* | ASP, GLU | 5.0 |
| Basic residues | name O* | ARG, LYS, HIS | 5.0 |
| Cys/Met residues | name S* C* | CYS, MET | 4.5 |
| Generic proximity | all ligand atoms | all atoms of candidate protein residues | 4.0 |

C*, N*, O*, H* and S* match atom names beginning with the indicated letter; H* includes hydrogens regardless of their bonded atom. Atom-name selections apply within each candidate protein residue. Ser/Thr and Asn/Gln classes were combined as polar residues for plotting. Classes overlap and describe distance-based proximity rather than chemically assigned interactions. Candidate residues were screened within 7 Å every fifth frame; the union was evaluated over all 5,001 frames.

**Table S5. Flexible receptor residues and docking scores of selected starting poses.**

| System | Flexible receptor residues | Selected pose | Selected FEB (kcal/mol) | Best FEB (kcal/mol) | Cluster size |
|---|---|---|---|---|---|
| HS+GBP | Trp205, Tyr236, Arg241, Trp243, Tyr450 | 3 | −8.057 | −8.313 | 6 / 100 |
| HS+PGB | Arg241, Tyr450, Asn550, Ser553, Gln554 | 2 | −7.309 | −7.413 | 18 / 100 |
| CE+GBP | Phe233, Thr238, Trp240, Tyr500, Asn568 | 8 | −6.054 | −7.147 | 5 / 100 |
| CE+PGB | Phe233, Thr238, Trp240, Tyr241 | 1 | −5.659 | −5.659 | 15 / 100 |

Scores are FEB values from AutoDockFR outputs. Cluster size indicates the number of docking solutions in the selected cluster out of 100 solutions. Starting poses were selected using docking scores, visual inspection and, in some cases, preliminary MD observations. Scores were compared within each search rather than as cross-system affinity estimates.

**Table S6**

#### A. Comparisons of structures associated with FEL minima

| System | Minima | Global backbone RMSD (Å) | Local backbone RMSD (Å) | Ligand RMSD after local fit (Å) |
|---|---|---|---|---|
| HS+GBP | 1-2 | 2.843 | 1.162 | 1.307 |
| HS+GBP | 1-3 | 4.999 | 1.833 | 5.699 |
| HS+GBP | 2-3 | 4.728 | 2.067 | 5.493 |
| HS+PGB | 1-2 | 3.680 | 1.060 | 0.548 |
| HS+PGB | 1-3 | 2.874 | 0.957 | 0.764 |
| HS+PGB | 2-3 | 3.400 | 0.793 | 0.526 |
| CE+GBP | 1-2 | 3.810 | 2.068 | 7.080 |
| CE+GBP | 1-3 | 3.007 | 1.198 | 4.984 |
| CE+GBP | 2-3 | 4.185 | 2.223 | 5.303 |
| CE+PGB | 1-2 | 7.041 | 1.781 | 4.240 |
| CE+PGB | 1-3 | 5.267 | 1.914 | 6.943 |
| CE+PGB | 2-3 | 4.447 | 1.507 | 3.821 |

Fits use equally weighted protein backbone atoms. Local selections are the within-system union of residues with any heavy atom within 8 Å of a ligand heavy atom in any of the three selected structures: 52 residues for HS+GBP, 40 for HS+PGB, 51 for CE+GBP and 37 for CE+PGB. Each system uses the same selection for all three snapshots. Ligands follow the local protein fit, with atoms matched by name and without independent ligand fitting or symmetry correction.

#### B. Snapshot times and relative grid energies

| System | **Minimum 1** Time (ns); relative energy (kJ/mol) | **Minimum 2** Time (ns); relative energy (kJ/mol) | **Minimum 3** Time (ns); relative energy (kJ/mol) |
|---|---|---|---|
| HS+GBP | 283.9 ns; 0.000 | 359.1 ns; 0.584 | 44.3 ns; 1.432 |
| HS+PGB | 424.1 ns; 0.000 | 92.1 ns; 0.226 | 210.5 ns; 0.266 |
| CE+GBP | 315.9 ns; 0.000 | 128.4 ns; 1.493 | 406.4 ns; 1.925 |
| CE+PGB | 208.8 ns; 0.000 | 482.9 ns; 0.303 | 349.7 ns; 1.886 |

Grid energies are in kJ/mol at 303.15 K, relative to the most occupied bin. Each selected frame is the frame nearest the selected bin centre in bin-width-normalized PC coordinates. A grid minimum is not an integrated basin or proof of an independent metastable state. Structural overlays are shown in Figure S2.

**Figure S1. MM/GBSA temporal correlation and block diagnostics.** Rows represent HS+GBP, HS+PGB, CE+GBP, and CE+PGB. Left: frame-wise estimates, overall means (dashed), and 50-ns means (black). Middle: normalized autocorrelation. Right: block-mean standard errors, with dashed lines indicating SD/√501. Equal blocking uses 500 frames. These diagnostics do not establish convergence or reliable confidence intervals. Numerical summaries are provided in Table S3.

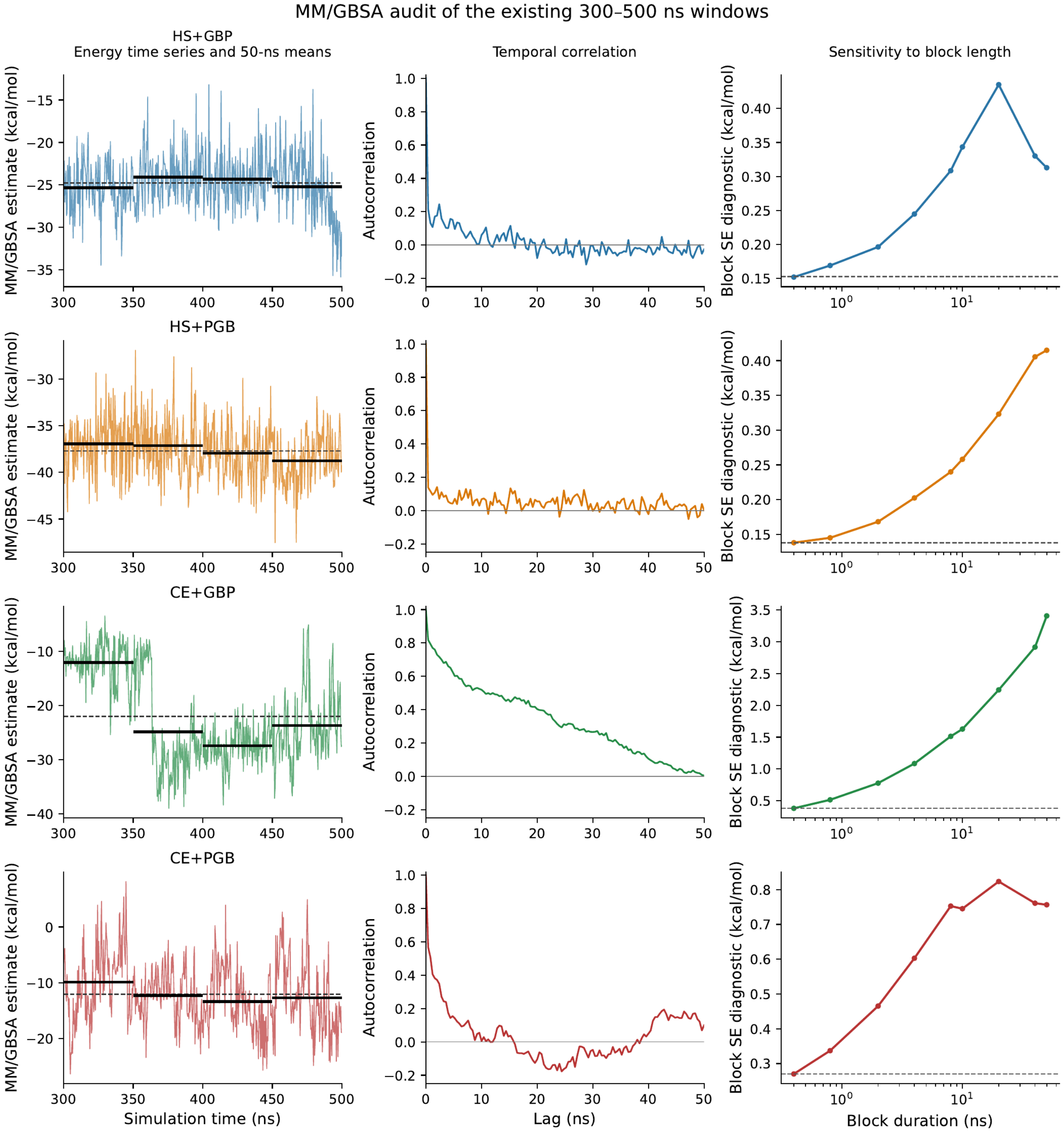

**Figure S2. Structural overlays of snapshots associated with selected FEL minima.** Left: whole-protein backbone alignment. Right: local backbone alignment using the residue selections defined in Table S6. Blue, orange, and cyan represent minima 1, 2, and 3, respectively, consistent with Figure 7. Ligands follow the protein alignment without independent fitting. Snapshot times are indicated for each system.

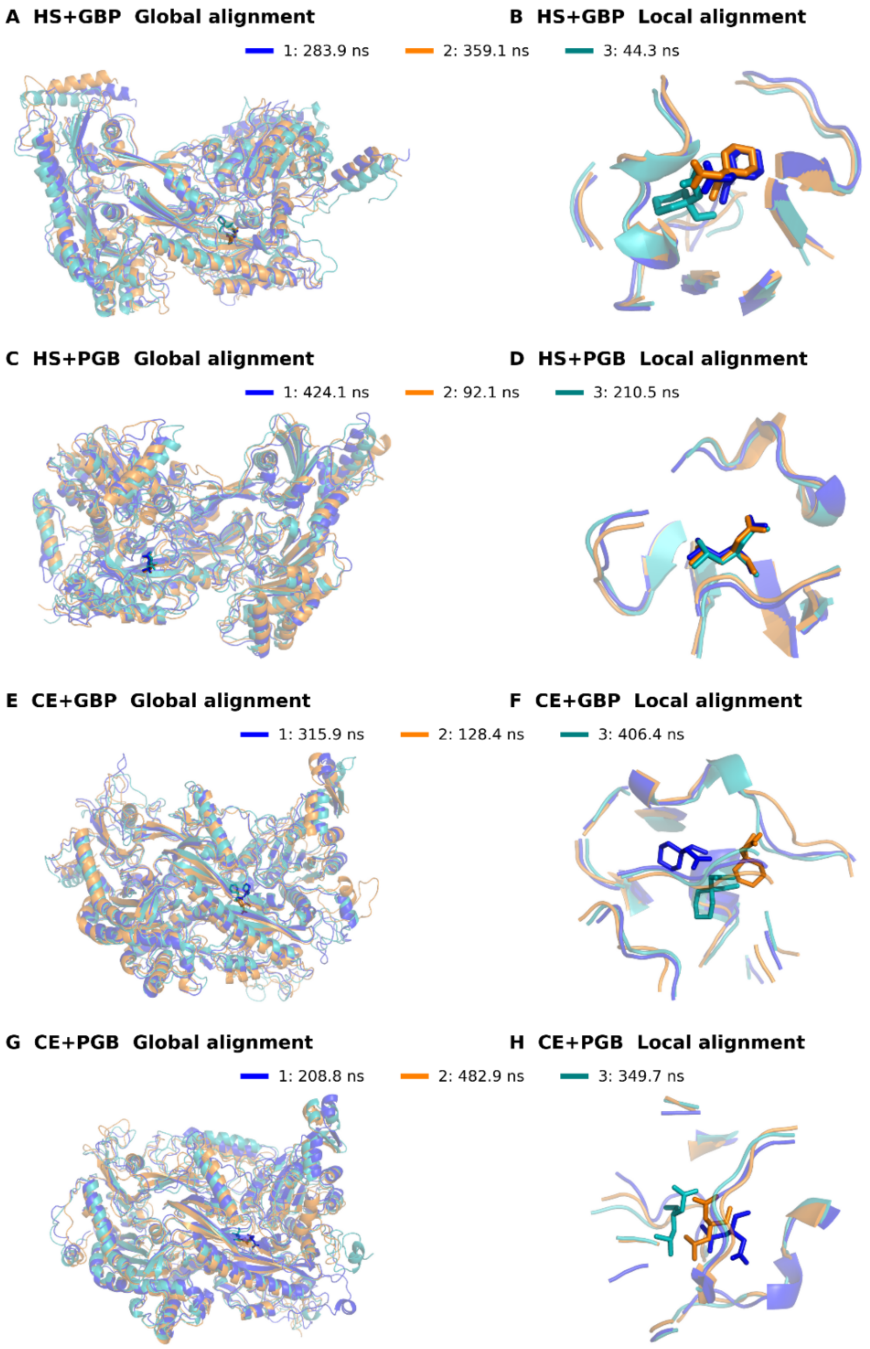
